# multiDEGGs: a multi-omic differential network analysis package for biomarker discovery and predictive modeling

**DOI:** 10.1101/2025.09.25.678475

**Authors:** Elisabetta Sciacca, Susan Wang, Costantino Pitzalis, Myles J Lewis

**Author notes:** Corresponding authors: Dr. Elisabetta Sciacca, Professor Myles J. Lewis, Professor Costantino Pitzalis. Co-senior last authors (Costantino Pitzalis, Myles J Lewis).

## Abstract

Modern clinical trials increasingly leverage high-throughput omic data for patient stratification and biomarker discovery. While traditional differential gene expression analysis disregards the networked nature of molecular entities and produces extensive gene lists with limited interpretability, differential network analysis has emerged as a crucial complementary analysis for comparative studies. Here we present multiDEGGs, a CRAN R package that enables differential network analysis in multi-omic scenarios.

multiDEGGs uses a multi-layer graph framework to model omic data by leveraging an internal network of over 10 000 literature-validated biological interactions. For each data type, differential networks are generated, and the statistical significance of each link (p-values or adjusted p-values) is evaluated through robust linear regression with interaction terms. These networks are then integrated into a comprehensive visualisation that allows interactive exploration of cross-omic patterns.

Beyond network visualization and exploration, multiDEGGs extends its utility to predictive modelling applications. The package facilitates seamless integration into cross-validation machine learning pipelines, serving as feature selection and augmentation tool.

We validated multiDEGGs using two cohorts of rheumatoid arthritis patients who underwent tocilizumab and rituximab therapy, respectively. For each treatment group, multi-layer differential interactions were identified, and seven machine learning models were trained to predict treatment resistance using synovial RNA-seq data. We systematically compared multiDEGGs against five traditional feature selection methods. On average, AUC values obtained with multiDEGGs showed an improvement of 0.10 compared to conventional filters.

**KEY POINTS:**

- Traditional gene expression analysis leaves researchers with hundreds of ‘significant’ genes but no clear biological story. The multiDEGGs CRAN package shifts the focus: instead of asking which genes change, it asks which gene *relationships* change.
- It can be used with single or multi-omic data: differential networks are calculated separately for each data type, with results integrated into a comprehensive, interactive view.
- multiDEGGs can be combined with the nestedcv CRAN package (nested cross-validation) to serve as feature selection and augmentation tool.
- In comparative evaluations, machine learning models trained with multiDEGGs-selected features showed AUC improvements of 0.10 compared to other feature selection methods.

## INTRODUCTION

Modern clinical trials increasingly leverage high-throughput omics data to characterize distinct patient phenotypes and elucidate differential biological mechanisms across various conditions. In this context, differential gene expression analysis (DGE) represents the most basic analytical approach, despite its limitations in efficacy and interpretability(Nance et al. 2022, Cui et al. 2021, Stretch et al. 2013, Ein-Dor and Domany 2006). Differential network analysis has emerged as a crucial complement to single-gene differential methods by identifying alterations in biomolecular relationships, thereby providing deeper insights into the molecular mechanisms underlying specific phenotypes or disease states. The past decade has witnessed significant advancements in the development of R packages dedicated to differential network analysis of high-throughput omics data. These tools can be broadly categorised based on their statical and mathematical strategies. One group of approaches primarily focuses on co-expression patterns derived directly from the omics data and identifying differential co-expressed modules. Methods in this category typically employ correlation measures, such as Pearson’s correlation coefficient, to construct networks and subsequently compare them between conditions. For example, the DiffCorr(Fukushima 2013) package calculates correlation matrices and employs Fisher’s z-test to identify differential correlations, while DiffCoEx(Tesson, Breitling and Jansen 2010) leverages the WGCNA framework to detect differentially co-expressed gene modules. WGCNA is also used – among other options – in the DCGL package(Liu et al. 2010), and the INDEED package(Li et al. 2018) allows users to choose between partial, Pearson, and Spearman correlation. In contrast, another class of tools integrates prior knowledge of biological interactions to guide the differential network analysis. NetOmics(Bodein et al. 2022), DIABLO(Singh et al. 2019) and DEGGs(Sciacca, Alaimo, Silluzio, Ferro, Latora, Pitzalis, Pulvirenti and Myles J. Lewis 2023) are examples of this approach. This strategy addresses the computational challenges of analysing all possible gene-gene combinations in NGS data while enhancing interpretability by focusing exclusively on functionally relevant interactions. Of those tools, DEGGs is the only one providing a user-friendly interactive interface, and differential statistical significance for each gene-gene link via robust linear regression with interaction term.

Here we present multiDEGGs (multi-omics Differentially Expressed Gene-Gene pairs)(Sciacca and Lewis 2025), an evolution of the DEGGs package that extends differential network analysis to multiomic scenarios using multi-layer differential networks (Video 1).

We applied multiDEGGs to two rheumatoid arthritis case studies to identify multi-layer differential networks that revealed key biological mechanisms distinguishing therapy responders from non-responders, offering valuable insights for clinical investigation.

In the second part of the paper, we also demonstrate how the resulting differential interactions can be effectively leveraged for feature selection and augmentation in machine learning applications. To this aim, the original code has undergone substantial speed and efficiency improvements from its previous version and has been expanded with new features. Finally, to enable seamless integration in nested cross-validation processes, we also extended the nestedCV package(Lewis et al. 2023, Myles Lewis et al. 2025) with complementary functionalities. The features selected by multiDEGGs led to better model performances when compared with other well-known filters (e.g. univariate filters, glmnet filter, ect.).

## MATERIALS AND METHODS

### Differential multi-omic analyses through multi-layer networks

The multiDEGGs package extracts differential multilayer networks following the workflow depicted in Figure 1.

**Figure 1.**
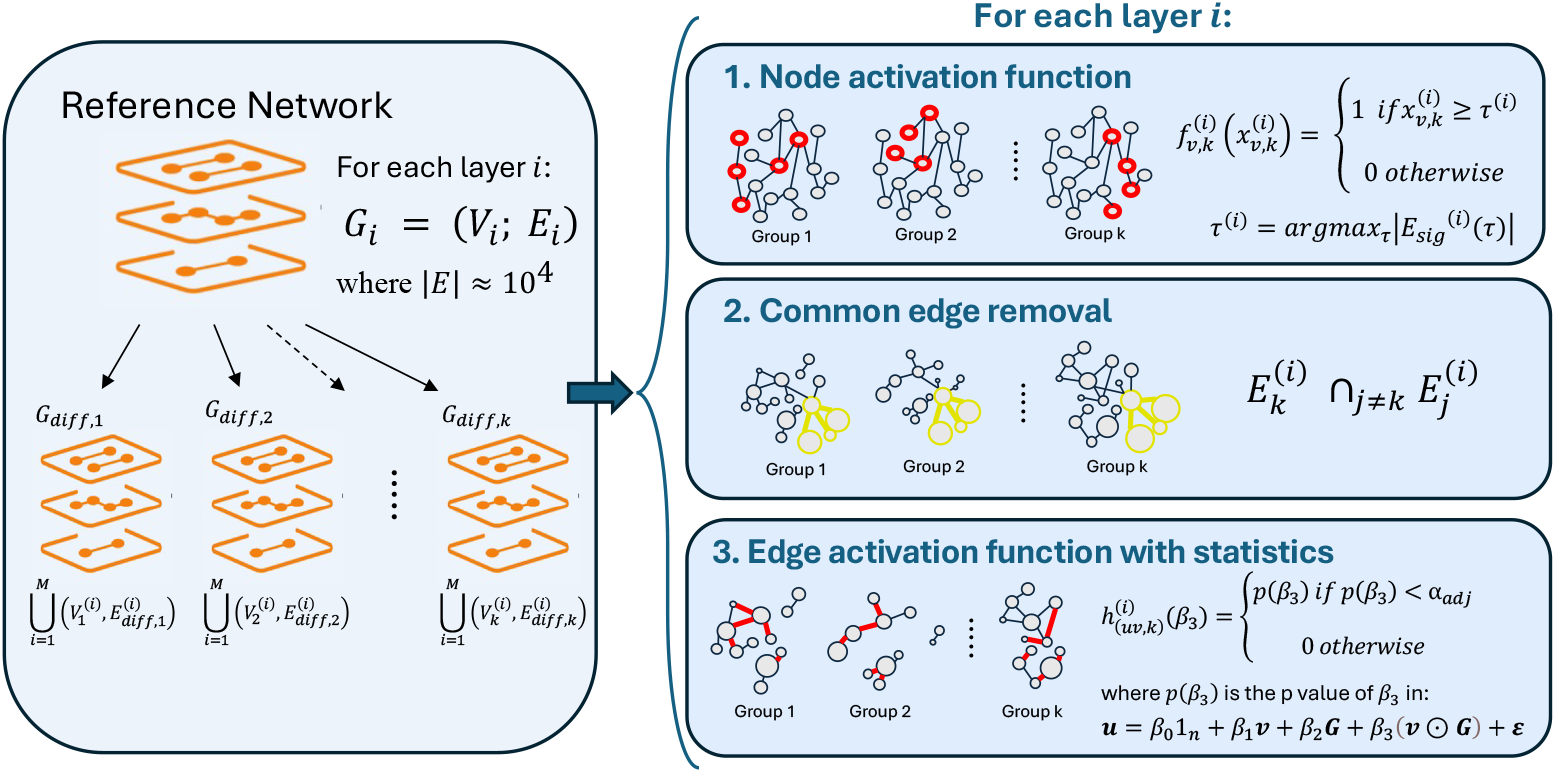
Workflow diagram showing the multiDEGGs internal implementation for multi-omic differential network analysis.

In particular, let

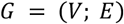

be a reference biological network with vertex set *v* representing molecular entities and edge set *E* representing known molecular interactions from literature, where |*E*| ≈10^4^. From this reference network, multiDEGGs extracts group-specific differential multilayer networks for each experimental group or condition *k*

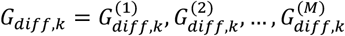

where each layer 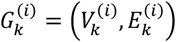represents a specific omic data type (e.g. RNA-seq, proteomics, Olink, etc.) filtered for group *k*. The key distinction between groups arises from the group-specific activation patterns of molecular entities.

During step 1, for each group *k* and layer *i*, multiDEGGs defines a node activation function 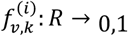 as a step function:

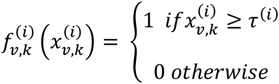

Here, 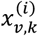 represents the normalised median expression or abundance of the biological entity *v* in group within the omic layer *i*, and τ^(*i*)^ is the activation threshold selected by solving:

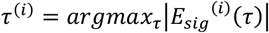

where 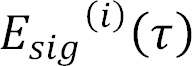 is the set of significant edges at threshold τ, as per percolation procedure previously detailed in (Sciacca, Alaimo, Silluzio, Ferro, Latora, Pitzalis, Pulvirenti and Myles J Lewis 2023), where the first version of the package was introduced. The group-specific active vertex set for layer *i* is then defined as

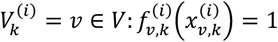

Since 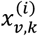 varies between groups, during the first step different nodes will be activated for each group *k*, creating group-specific network topologies. To retain only the group-specific edges, in step 2 multiDEGGs identifies and removes edges that are common to all *k* groups within each omic layer. For layer *i*, the group-specific edge set is then defined as

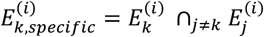

where 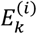 represents the edges connecting active nodes in group *k* and layer *i*. In step 3, an edge activation function is applied to each remaining group-specific edge in layer *i*. multiDEGGs employs robust linear regression with interaction term to assess whether the differential connectivity between groups is statistically significant. For each potential edge (*u, v*) connecting active nodes within the same layer *i* and group *k*, let ***u, v*** ∈ *R*^*n*^ be the corresponding normalised expression or abundance vectors of the two biological entities across all samples, and ***G*** ∈ 1,2, …, *K*^n^ be the categorical group vector indicating group membership for each sample. multiDEGGs formulates the regression model as:

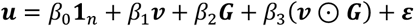

where **1**_*n*_ is the vector of ones of dimension *n*, ⊙ denotes the element-wise (Hadamard) product, *β*_3_ captures the differential interaction effect between the two biological entities in group *k* relative to other groups, and ***ε*** ∈ *R*^*n*^ represents the error term vector. The differential edge activation function 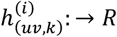 is then defined as

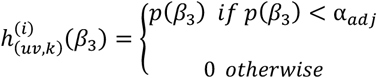

where *p*(*β*_3_) is the adjusted p value for the interaction term *β*_3_, and α_*adj*_ is the multiple testing corrected significance threshold defined by the user. The differential edge set for layer *i* is then defined as:

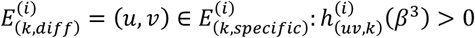

Therefore, the complete group-specific differential multilayer network is represented as:

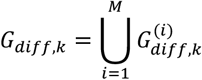

where 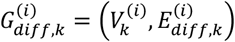 is the differential graph for group *k* and layer *i*.

It is worth noting that, in this final representation, each molecular entity is active in only a subset of layers. Being able to visually identify those entities active in multiple omic layers – forming bridging connections – is important to shed light on possible cross-layer signal propagation associated to certain conditions.

For visualization purposes, the final multi-layer differential networks are integrated into a comprehensive display where each omic layer *i* is assigned a distinct colour, enabling users to easily gain an overview of all layers and explore cross-omic patterns. The interactive visualization is implemented through the `View_diffNetworks()` function using R Shiny and includes several features: a slider for filtering network links by p-value (or adjusted p-value), a search box to locate specific genes or molecular entities, clickable links that display the associated differential regression plot when selected, and clickable nodes that generate boxplots showing expression differences between experimental groups (Video 1).

Overall, this framework ensures group-specific differential network extraction from a common reference network, layer-specific activation through percolation-optimised thresholds, removal of common connectivity to highlight group-specific patterns, differential edge weighting based on the continuous significance values of interaction effects within each layer, and robustness through multiple testing correction.

## RESULTS

### Multi-omic differential network analysis in rheumatoid arthritis

We applied multiDEGGs to two rheumatoid arthritis multi-omic datasets. Rheumatoid arthritis (RA) is a chronic autoimmune disease characterized by synovial inflammation and joint destruction. Despite the availability of diverse biologic disease modifying anti-rheumatic drugs targeting distinct immune pathways, 30–40% of RA patients exhibit inadequate response (Taylor et al. 2022), and approximately 10% develop multi-refractory disease, failing more than two biologic classes in longitudinal studies (Novella-Navarro et al. 2020). To better understand the biological signatures underlying treatment response, multiDEGGs has been applied to two cohorts of rheumatoid arthritis patients who underwent tocilizumab and rituximab therapy, respectively (Rivellese et al. 2022, Humby et al. 2021, Myles J. Lewis et al. 2025, Rivellese et al. 2023). In both studies, synovial biopsies were collected before treatment initiation, and therapeutic response was assessed following treatment completion. The resulting multi-layer differential networks highlighted important mechanisms distinguishing patients who respond to therapy from those who do not, providing crucial insights for clinical investigation.

### Multi-omic differential network analysis in RA patients not responding to tocilizumab therapy

The RA cohort of patients treated with tocilizumab consisted of 65 individuals with available synovial RNAseq (n=65), O-link data (n=64), and lower numbers of mass spectrometry proteomic (n=17) and phosphoproteomic (n=17) data.

The main cluster of the resulting multi-omic differential network centres on *PIK3R1*, which shows extensive synovial RNA-seq co-expression (green arrows) with key cytokine-driven inflammatory pathway signalling molecules including *JAK1, JAK3, TYK2, IL2RG, CSF2RB*, and *AKT2* (Figure 2). This indicates possible coordinated activation of the PI3K–AKT and JAK/STAT pathways, both of which are downstream of cytokine receptors, including the IL-6 receptor (IL-6R) (Malemud 2018). The co-expression of *ADCY4* and *PRKACA* (Padj = 0.0074) in responders suggests coordinated activation of the cAMP–PKA pathway. *ADCY4* encodes adenylate cyclase 4, which generates cAMP as a second messenger, while *PRKACA* encodes the catalytic subunit of protein kinase A, the primary effector of cAMP signalling. Together, their co-expression implies enhanced cAMP-driven phosphorylation of downstream targets, which is known to increase secretion of cytokines including IL-6 (Hershko et al. 2002). Phosphoproteomic links (orange arrows) involving ITGB2 and ICAM3 indicate active integrin-mediated adhesion and immune cell retention, which may reinforce cytokine-driven inflammation.

**Figure 2.**
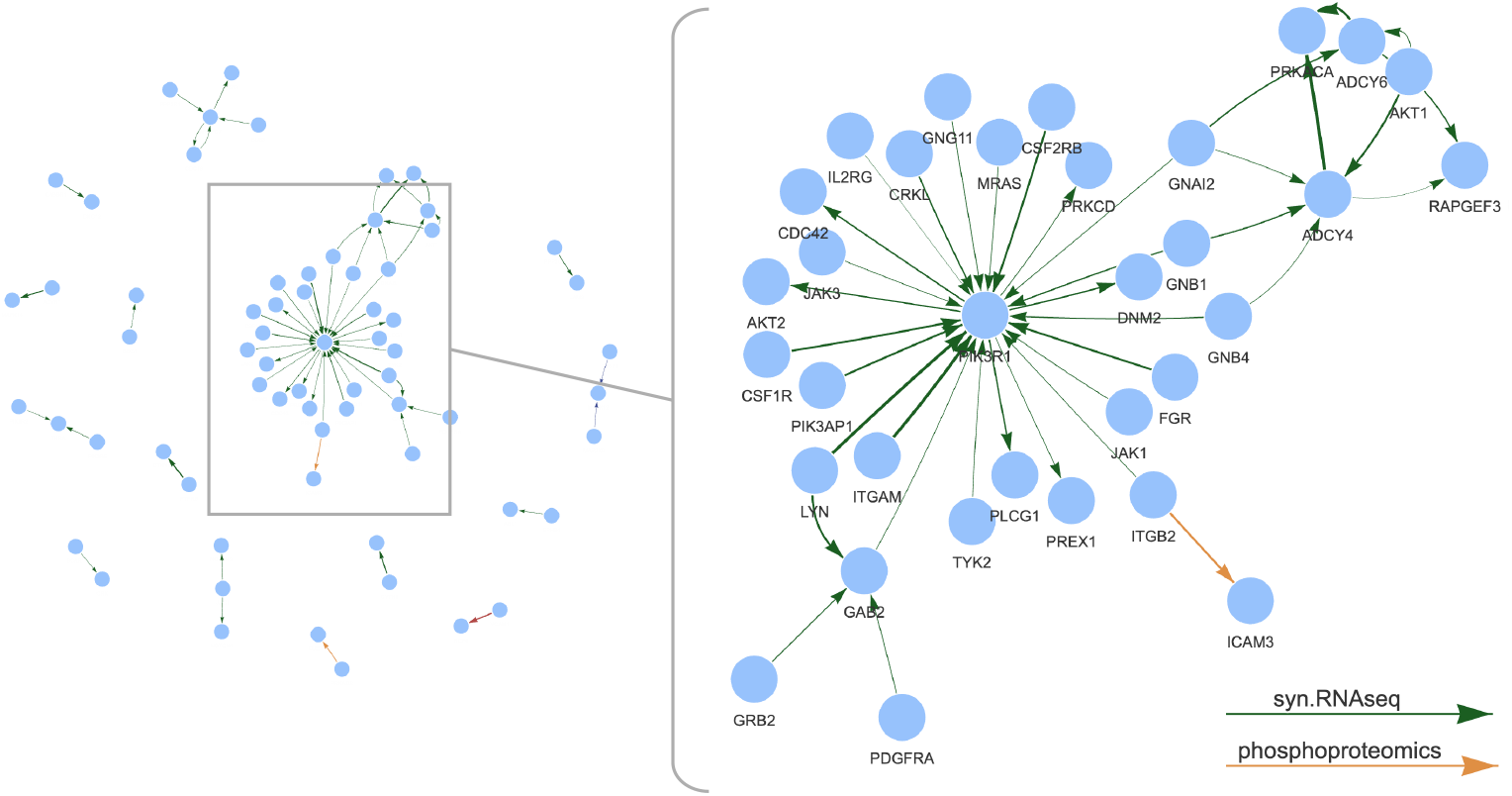
Network overview and magnified view of a cluster of multi-omic differential interactions between tocilizumab responders and non-responders in rheumatoid arthritis patients. The network displays differential interactions with edge colours representing different omics data types (green edges for synovial RNA-seq, orange edges for phosphoproteomics data). All interactions shown have adjusted p-values < 0.05. Edge thickness is proportional to statistical significance, with thicker edges representing lower adjusted p-values. Arrow directions reflect literature-reported regulatory relationships and are not derived from data.

From a clinical perspective, the presence of this integrated network prior to therapy suggests that the pathology in responders is highly dependent on cytokine, including IL-6, mediated PI3K–AKT and JAK/STAT signalling for synovial inflammation. Tocilizumab, by blocking IL-6R signalling, likely disrupts this central network, leading to a more profound therapeutic effect in these patients. Conversely, the absence of this transcriptional and phosphoproteomic configuration in non-responders suggests that their synovial inflammation is driven by alternative, IL-6-independent pathways.

### Dysregulated multi-omic interactions in RA patients treated with rituximab

The RA cohort of patients receiving rituximab treatment comprised 72 patients with accessible synovial RNAseq and O-link data (n=72) along with fewer mass spectrometry-based proteomic and phosphoproteomic (n=21) profiles.

The resulting multi-omic differential network highlights *ITGB3* and *ITGA3* as central genes (Figure 3). They encode integrin subunits (β3 and α3) that mediate cell-extracelluar matrix (ECM) adhesion and signalling, processes central to synovial fibroblast activation and immune cell trafficking in RA (Lowin and Straub 2011).

**Figure 3.**
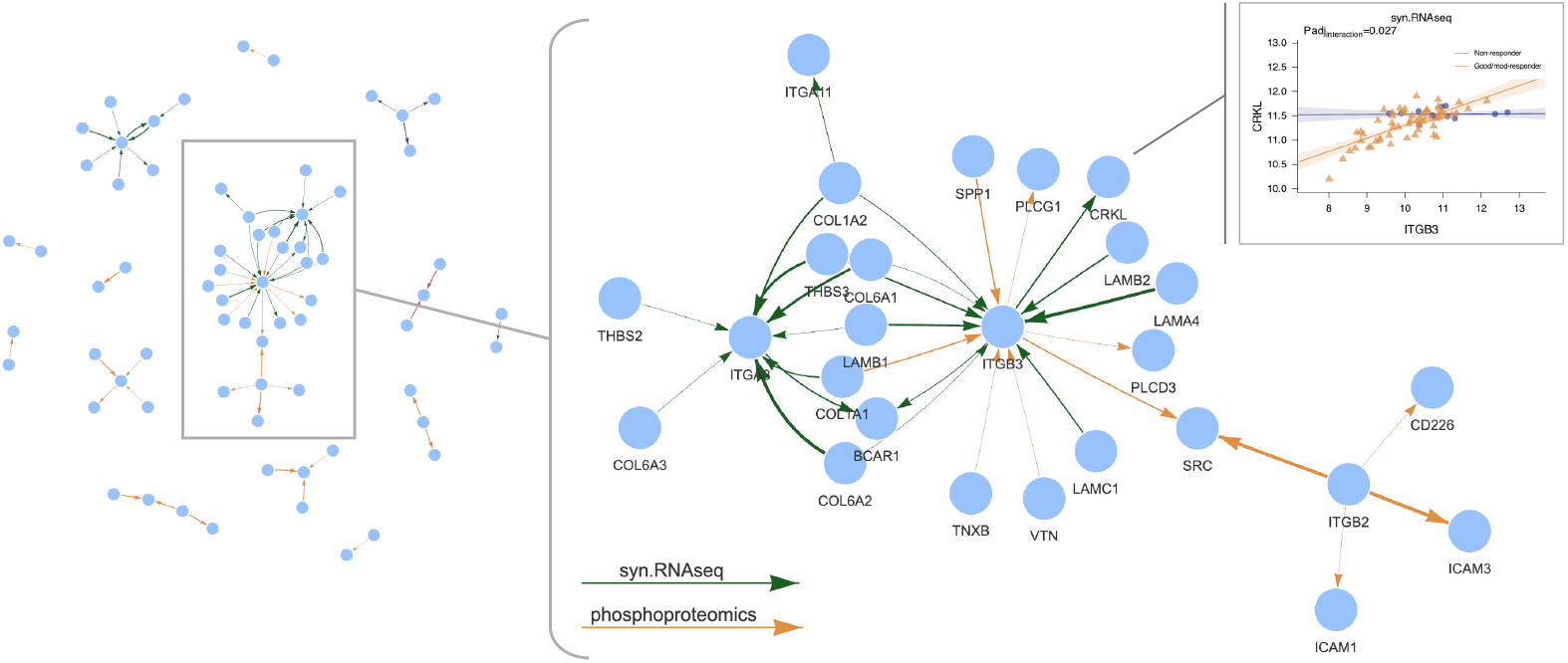
Network overview with magnified view of a cluster of multi-omic differential interactions between rituximab responders and non-responders in rheumatoid arthritis patients. The network displays differential interactions with edge colours representing different omics data types (green edges for synovial RNA-seq, orange edges for phosphoproteomics data). All interactions shown have adjusted p-values < 0.05. Edge thickness is proportional to statistical significance, with thicker edges representing lower adjusted p-values. Arrow directions reflect literature-reported regulatory relationships and are not derived from data.

Their presence suggests integrin-mediated (ECM) interactions, which represent a key distinguishing factor for rituximab responders in RA. *ITGB3* expression is strongly associated with multiple extracellular matrix component genes (*LAMA4, LAMB1, LAMB2, COL1A1, COL6A2, THBS3, SPP1*) in responders, consistent with its role in modulating fibroblast adhesion, migration, and tissue remodelling in the synovium. Notably, the association of ITGB3 with SRC and PLCG1 through phosphoproteomics data suggests downstream activation of focal adhesion and MAPK signalling pathways, which regulate pro-inflammatory and survival signals. Through SRC, this phosphoproteomic network extends to ITGB2, a leukocyte-specific integrin central to immune cell adhesion and migration (Lowin and Straub 2011). The ITGB3–SRC–ITGB2 axis may facilitate immune cell, particularly B cell, recruitment into the inflamed synovium, aligning with the higher B cell burden typically observed in rituximab responders (Rivellese et al. 2022). The upper right plot demonstrates a significant positive correlation between *ITGB3* and *CRKL* (an adaptor protein involved in integrin and growth factor signalling) expression in good/moderate responders but absent in non-responders. The strong coupling of *ITGB3–CRKL* expression in responders indicates that integrin-mediated signalling dynamics, potentially linked to cellular adhesion and immune cell trafficking, are more transcriptionally coordinated in patients who benefit from rituximab.

The ITGA3 network also shows strong co-expression with ECM-related genes, including *COL1A1, COL1A2, COL6A1, COL6A2, COL6A3, LAMB1, THBS2*, and *THBS3*, indicating active matrix remodelling and cell–matrix adhesion in rituximab responders. This pattern suggests that fibroblast-like synoviocytes are engaged in strong integrin–ECM interactions, promoting a structural microenvironment conducive to immune cell infiltration and retention.

Collectively, these networks may indicate that responders possess a synovial microenvironment preconditioned for immune cell (including B cell) migration, retention, and survival via integrin– ECM interactions. As rituximab targets CD20+ B cells, its efficacy is enhanced in B cell–rich synovium, which may explain why responders display an *ITGB3/ITGA3*-centred network of transcriptional and phosphoproteomic associations that is absent in non-responders.

## MACHINE LEARNING APPLICATIONS

### multiDEGGs as feature selection and feature augmentation method in machine learning

DEGGs(Sciacca, Alaimo, Silluzio, Ferro, Latora, Pitzalis, Pulvirenti and Myles J. Lewis 2023) was initially conceived as differential network analysis tool, with a focus on the interactive exploration of networks to allow easy and quick identification of important nodes, clusters and biological interactions. However, beyond discovery, the identified key differential interactions between biomolecules can subsequently be utilised to predict clinical outcomes in new patients. In the context of machine learning with high-throughput data, researchers frequently encounter scenarios where the number of predictors (P) substantially exceeds the sample size (s). This dimensional imbalance requires selective feature inclusion, both for mathematical and clinical reasons. Models trained in P >> s conditions are prone to overfitting, resulting in high variance, poor generalisability, and spurious correlations that lead to mathematically unreliable predictions with inflated performances on trained data but poor real-world accuracy. Secondary, the use of all variables available in high-throughput data would not be feasible in clinical practise beyond research studies. For example, the implementation of targeted panels with selected, predictive biomarkers represents a more feasible approach than collecting whole transcriptome RNA sequencing in clinical routine.

Traditional feature selection approaches focus on identifying individual predictors that demonstrate the strongest association with the outcome of interest (e.g. univariate filters such as t-test or Wilcoxon test). However, since biological systems operate through complex networks of interacting components, high predictive power may also reside in feature combinations and interactions. In general, feature engineering involves a set of techniques that enables the creation of new features by combining or transforming the existing ones(Zheng and Casari 2018, Nargesian et al. 2017). For example, interaction terms are usually captured by multiplying or dividing two or more original features, while polynomial features extend this concept to include powers of individual variables and their combinations.

Such a mathematical framework allows models to capture complex, non-additive relationships where the effect of one variable depends on the value of another. The use of combined predictors and polynomial transformations thus offers significant potential to enhance machine learning models because they enable the integration of non-linear biological mechanisms that individual features fail to represent.

The higher-order informative content carried by differential interactions, combined with the multiDEGGs’ ability to extract only differential links validated in literature, positions it as particularly suitable both for single feature selection and for guiding the creation of modified predictors in machine learning applications, where conventional black-box algorithms may select single predictors lacking biological significance, thereby compromising the credibility and explainability of the resulting model in clinical settings.

DEGGs has already demonstrated potential in identifying gene-gene interactions predictive of treatment response in rheumatoid arthritis(Sciacca et al. 2022), however the validation of its efficacy as both a feature selection method and a predictor modification strategy requires evaluation in larger cohorts with rigorous cross-validation protocols. Previous research has shown that applying filtering across the entire dataset introduces bias when assessing model accuracy(Vabalas et al. 2019). Feature selection and predictor modification should be conducted exclusively on training data within cross-validation loops to prevent information leakage from the test set. To address this methodological requirement, we have expanded the nestedCV R package(Lewis et al. 2023) with a new functionality that enables the nested modification of predictors within each outer fold, ensuring that the attributes learned from the training part are applied to the test data without prior knowledge of the test data itself (Figure 4). The selected and combined features, and corresponding model, can then be evaluated on the hold-out test data without introducing bias.

**Figure 4.**
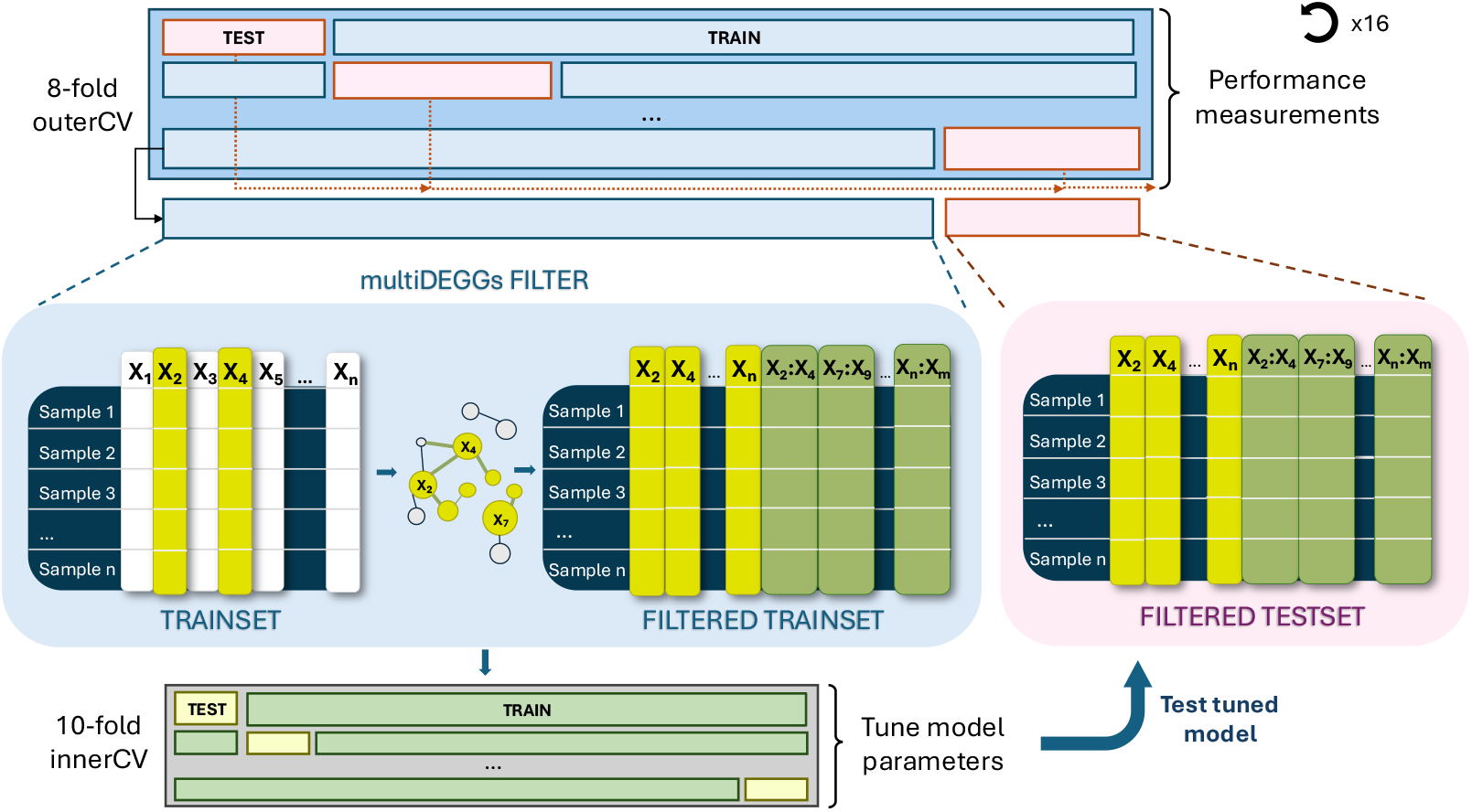
Nested cross-validation machine learning framework using multiDEGGs-based feature engineering.

Specifically, the nestedCV code has been extended to accept any user-defined function that filters or transforms the feature matrix by passing the function name to the `modifyX` parameter of the `nestcv.train()` function. A boolean flag (`modifyX_useY`) determines how the transformation is applied. When set to `TRUE`, the transformation is fitted using both the training response labels (i.e. the dependent variable) and the training feature matrix. The fitted model is then stored and used by another user-defined `predict()` function to modify the feature space in both training and testing data (Figure 5). This approach is suitable for supervised transformations that need to learn from the training data, such as PCA or the differential network analysis discussed in this paper. For unsupervised transformations like scaling, normalization, or polynomial feature generation that do not require knowledge of the response variable, `modifyX_useY` must be set to `FALSE`.

**Figure 5.**
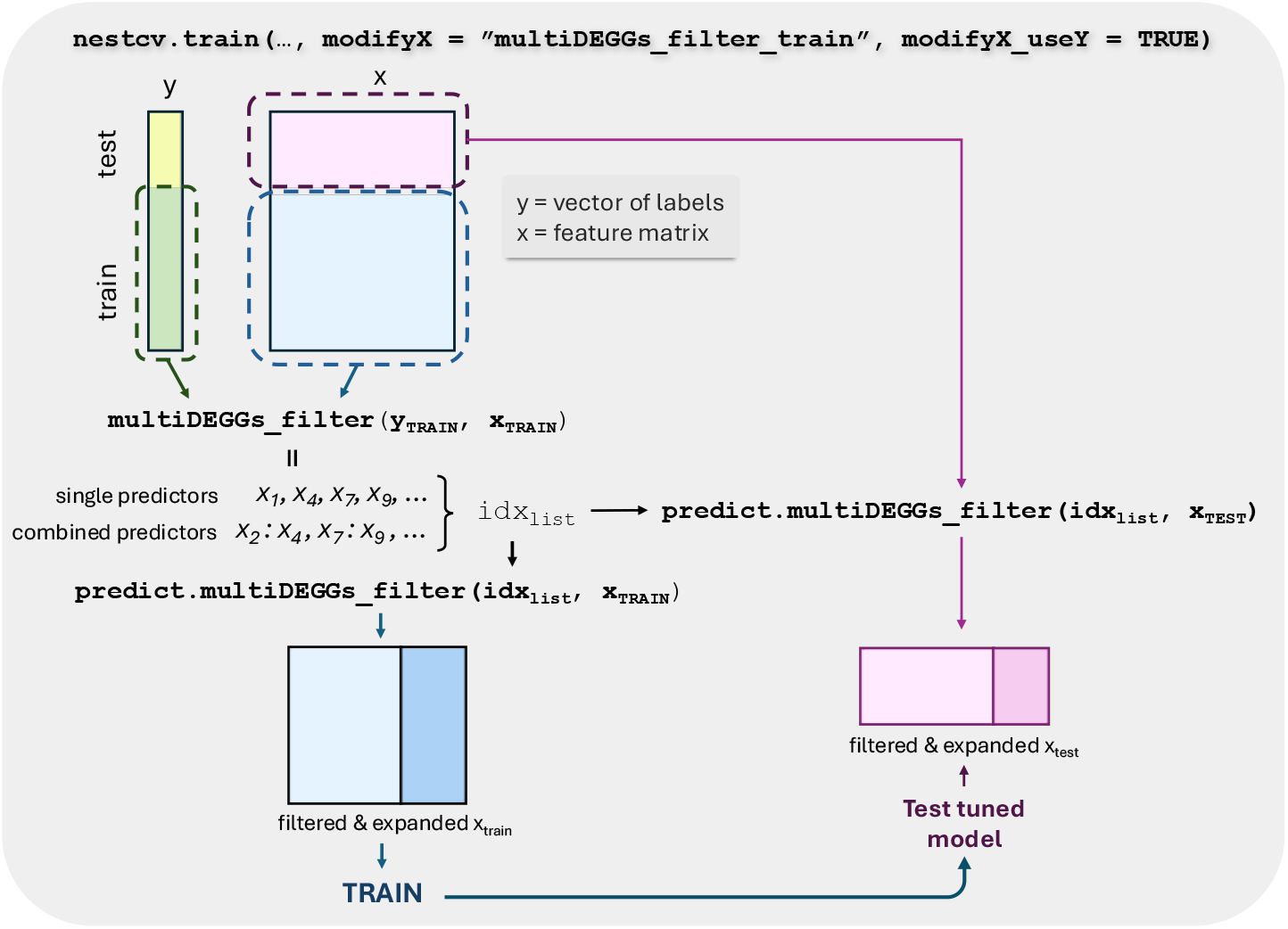
Code architecture for integrating multiDEGGs-based feature processing with the nestedcv cross-validation framework.

For this implementation, we then defined – and included in the multiDEGGs package – the filtering and predict functions as follows.

The `multiDEGGs_filter()`function identifies the differential molecular interactions, extracts both individual and paired corresponding variables, and stores them into an S3 model object containing the indices of each selected feature.

The second function (`predict.multiDEGGs_filter()`) is the predict method that applies the learned transformation to new data. It takes the paired variables identified during training and creates ratio features (feat. A / feat. B), while also retaining the individual predictors involved in the differential interaction. This ensures that both training and test data of each outer fold undergo the same feature transformation based on the patterns learned from the training set (Figure 5).

Finally, it is also worth noting that while traditional filtering functions require the number of selected features to be set by the user, multiDEGGs automatically establishes the best number of predictors thanks to the internal percolation process. Therefore, although a maximum threshold can still be set to prevent excessive feature inclusion, the exact number of selected features is automatically fine-tuned, eliminating arbitrary decision-making.

### Comparing multiDEGGs with other traditional filters

To evaluate the impact of multiDEGGs feature selection on machine learning performance, we trained seven machine learning models to predict treatment resistance using synovial RNA-seq data from the two aforementioned cohorts. For each model, we systematically compared multiDEGGs against five traditional feature selection methods: partial least squares (PLS), t-test, Wilcoxon rank-sum test, random forest, and regularized linear models (glmnet). To ensure fair comparison, we set the same maximum number of final features for all six filtering approaches (40 for the tocilizumab cohort and 50 for the larger rituximab cohort).

Finally, to guarantee robust evaluation and prevent data leakage, each model/filter combination has been trained 16 times using nested cross-validation with 8 outer folds and 10 inner folds (Figure 4). Figure 6 and 7 report the area under the receiver operating characteristic curve (AUC) obtained for each combination of model and filter.

**Figure 6.**
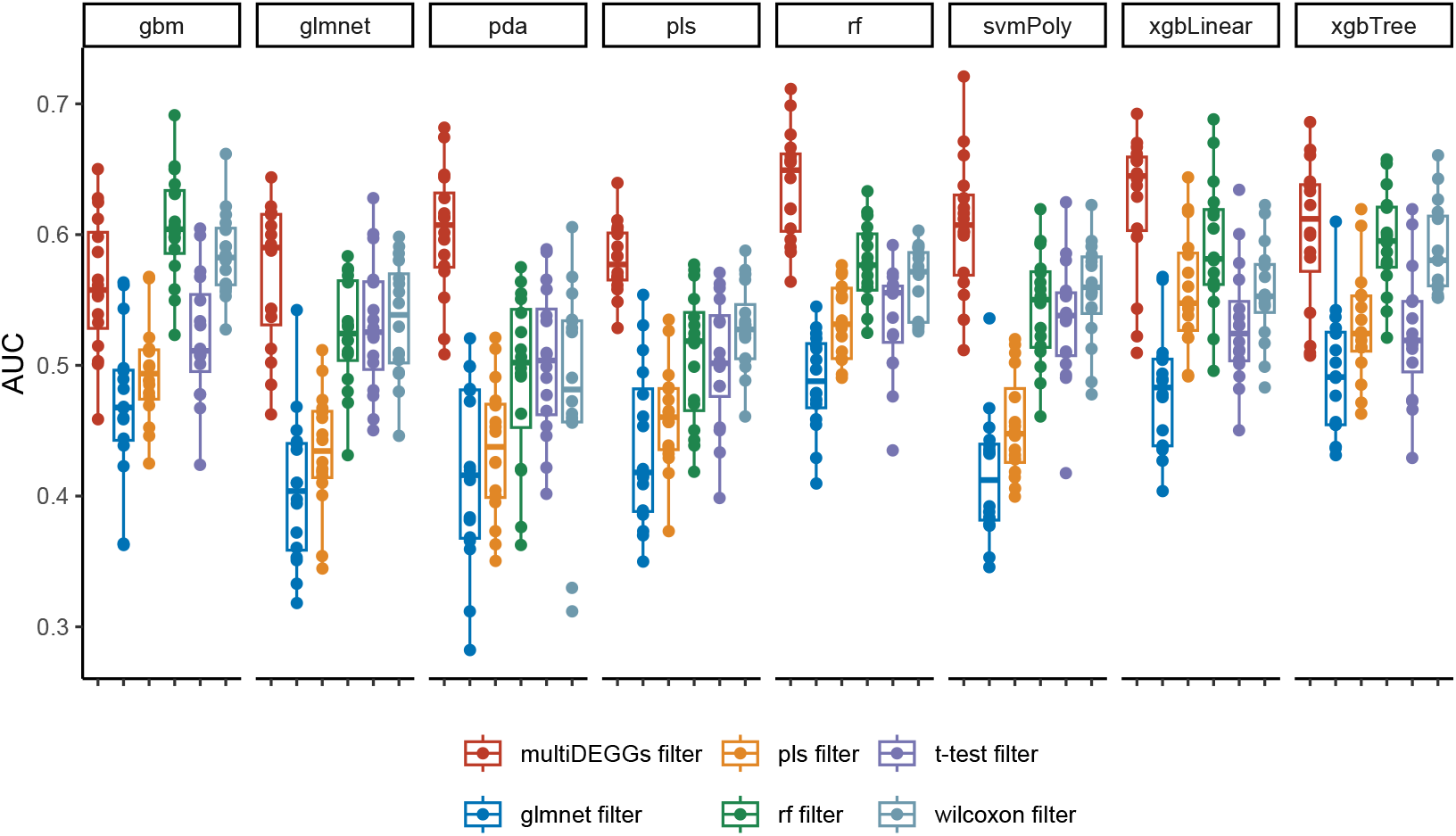
Boxplots showing AUC values of each trained model in the prediction of the tocilizumab resistant state of rheumatoid arthritis patients. Seven different models were tested: Generalized Linear Model with Elastic Net Regularization (glmnet), Penalized Discriminant Analysis (pda), Partial Least Squares (pls), Random Forest (rf), Support Vector Machine with Polynomial Kernel (svmPoly), Extreme Gradient Boosting with Linear Booster (xgbLinear), and Extreme Gradient Boosting with Tree Booster (xgbTree). For each model, multiDEGGs was sistematically compared against five traditional filtering methods: pls, T-test, glmnet, rf, and Wilcoxon filter. The maximum number of selected features was set to 40 for all filters.

**Figure 7.**
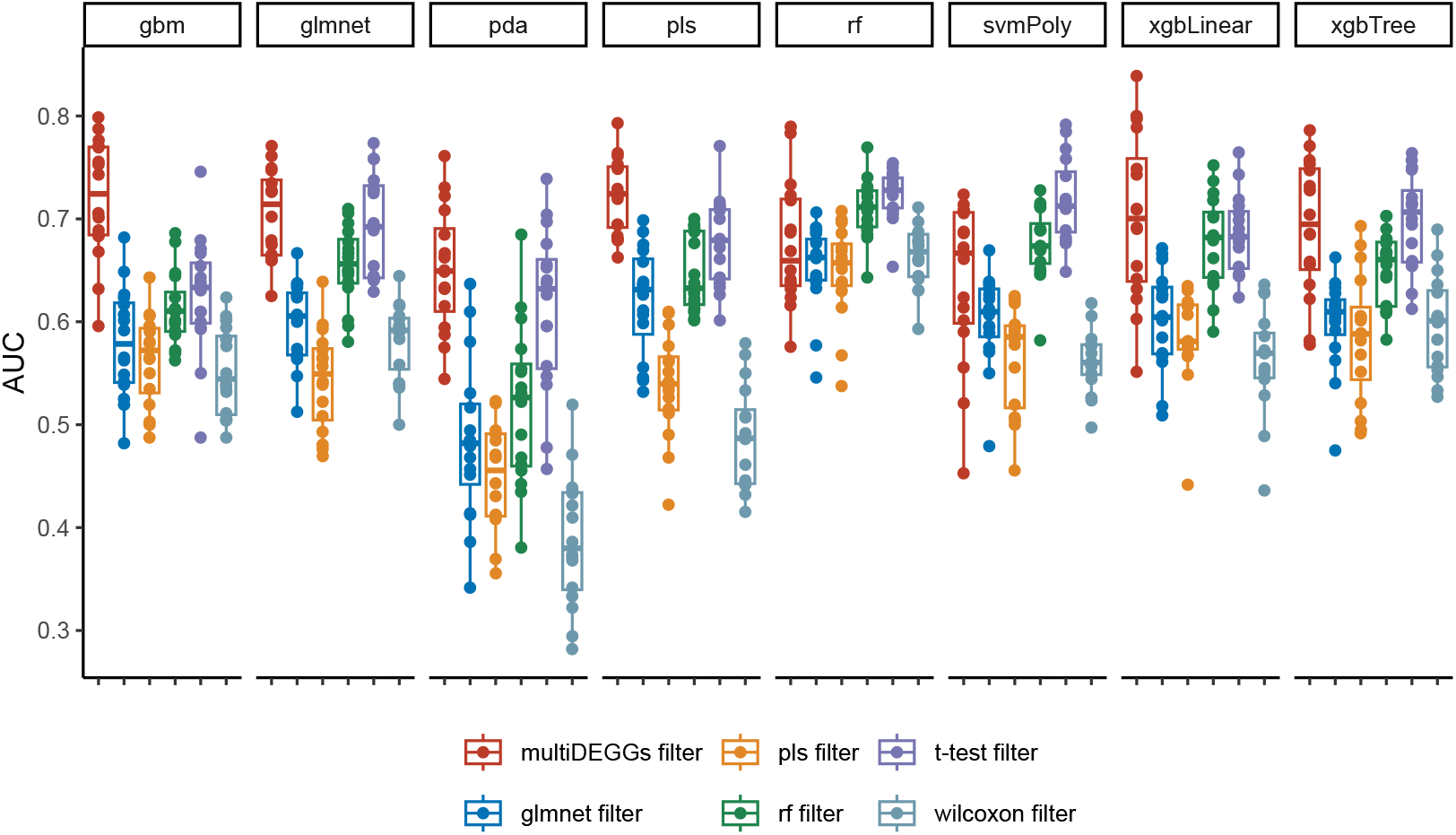
Boxplots showing AUC values of each trained model in the prediction of the rituximab resistance for rheumatoid arthritis patients. Seven different models were tested: Generalized Linear Model with Elastic Net Regularization (glmnet), Penalized Discriminant Analysis (pda), Partial Least Squares (pls), Random Forest (rf), Support Vector Machine with Polynomial Kernel (svmPoly), Extreme Gradient Boosting with Linear Booster (xgbLinear), and Extreme Gradient Boosting with Tree Booster (xgbTree). For each model, multiDEGGs was sistematically compared against five traditional filtering methods: pls, T-test, glmnet, rf, and Wilcoxon filter. The maximum number of selected features was set to 50 for all filters.

On average, starting from 22’975 protein coding genes available in synovial RNAseq, multiDEGGs reduced the feature space to 29 predictors for the tocilizumab cohort and 30 features for the rituximab cohort. Model performances obtained with multiDEGGs consistently outperformed the models trained with the alternative feature selection methods, particularly when using random forest and support vector machines. On average, AUC values obtained with multiDEGGs incremented by 0.10. To ensure this performance improvement is not inflated due to the presence of predictors as both single and combined variables, we repeated the same analysis by setting the `keep_single_genes` parameter of the `predict.multiDEGGs()`function to `FALSE`. This ensures that only combined predictors derived from differential pairs are selected as features, and the list of single variables stored by `multiDEGGs_filter_train()`function is ignored. As the feature space considerably decrease in this case, the filtering threshold was set to 15 for the tocilizumab cohort and to 25 for the larger rituximab group. Although its overall power decreased as a consequence of the smaller feature space, multiDEGGs confirmed better feature selection in most models (Supplementary Figure 1 and 2). In particular, in the tocilizumab cohort models trained with multiDEGGs obtained higher AUC values in five models out of eight, while the random forest filter’s AUC values were comparable or slightly higher for the remaining models. On the other hand, the rituximab dataset showed higher performances in four models out of eight, but performed poorly for the remaining models (Random Forest, Support Vector Machine with Polynomial Kernel, Extreme Gradient Boosting with Linear Booster, and Extreme Gradient Boosting with Tree Booster), indicating higher sensitivity to feature space reduction and insufficient filter power with fewer features.

## OTHER TECHNICAL CONSIDERATIONS

### Package Dependency Analysis and Optimization

The complexity of package dependencies has emerged as a critical consideration in R package development, particularly within the bioinformatics ecosystem where numerous interdependent packages can significantly impact installation time, maintenance burden, and computational resource requirements(Gu and Hübschmann 2022). Dependency heaviness, defined as the number of additional packages that a parent uniquely brings to a child package, provides a quantitative framework for assessing and optimizing package dependency structures. To evaluate and minimize the dependency burden of our package, we employed the pkgndep (v1.99.3), which systematically analyzes all packages listed in the Depends, Imports, LinkingTo, and Suggests/Enhances fields of the DESCRIPTION file. The initial dependency analysis performed on the original DEGGs code revealed the complete dependency tree with 86 packages directly required for installation (Supplementary Figure 2). For the development of multiDEGGs, we identified potential optimization targets and reduced the dependency heaviness by replacing heavy parent packages with lighter alternatives and removing non-essential dependencies, resulting in a substantially streamlined package architecture with 54 packages directly required (Supplementary Figure 3). This optimization process not only reduces the installation footprint and potential for dependency conflicts but also enhances the package’s long-term maintainability and accessibility for end users.

## CONCLUSIONS

multiDEGGs performs multi-omic differential network analysis by revealing differential interactions between molecular entities (genes, proteins, transcription factors, or other biomolecules). For each omic dataset provided, a differential network is constructed where links represent statistically significant differential interactions between entities. These networks are then integrated into a comprehensive visualisation that allows interactive exploration of cross-omic patterns, such as differential interactions present at both transcript and protein levels. For each link, users can access differential statistical significance metrics (p values or adjusted p values) and differential regression plots.

multiDEGGs can also be used as feature selection/augmentation tool in machine learning pipelines. Models trained with features engineered my multiDEGGs constantly improved their performances compared to traditional filters.

## DATA AVAILABILITY

All RNA-Seq data used in this article is available in <u>ArrayExpress</u>, and can be accessed with accession ID E-MTAB-13733 and E-MTAB-11611.

## ACKNOWLEDGEMENTS

The authors acknowledge the investigators and participants of the STRAP trial jointly funded by the UK Medical Research Council (MRC) and Versus Arthritis (grant no. MR/K015346/1); and the R4RA trial funded by the UK National Institute for Health and Care Research (NIHR) Efficacy and Mechanism Evaluation (EME) Programme (grant no. 11/100/76) for supporting this study and making their data available for this research. This work also acknowledges the support of the NIHR Barts Biomedical Research Centre (NIHR203330).

