## Supplementary Figure 2 for "multiDEGGs: a multi-omic differential network analysis package for biomarker discovery and predictive modeling"

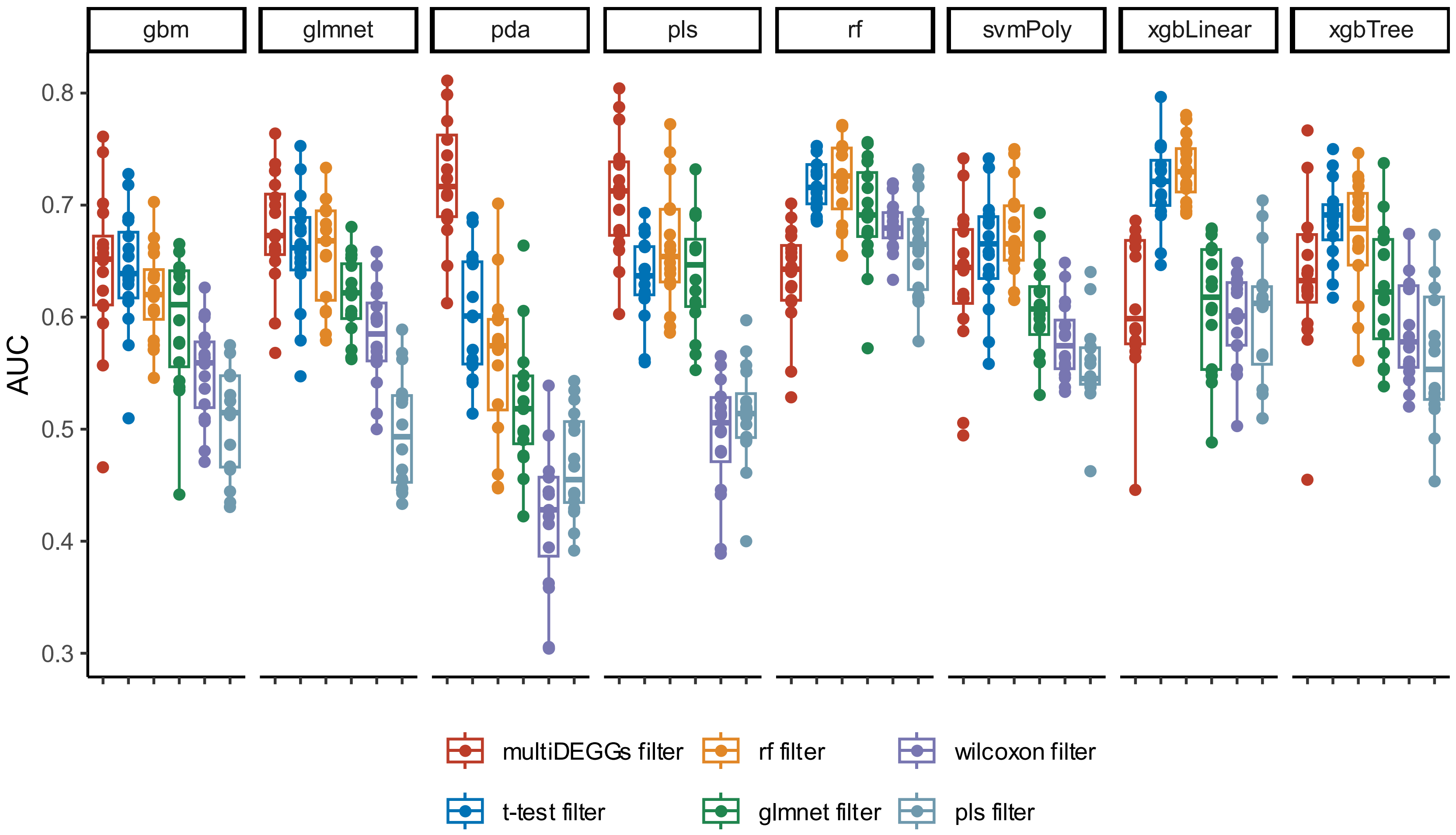


Supplementary Figure 2 – Boxplots showing AUC values of each trained model in the prediction of the rituximab resistant state of rheumatoid arthritis patients. Seven different models were tested: Generalized Linear Model with Elastic Net Regularization (glmnet), Penalized Discriminant Analysis (pda), Partial Least Squares (pls), Random Forest (rf), Support Vector Machine with Polynomial Kernel (svmPoly), Extreme Gradient Boosting with Linear Booster (xgbLinear), and Extreme Gradient Boosting with Tree Booster (xgbTree). For each model, multiDEGGs was sistematically compared against five traditional filtering methods: pls, T-test, glmnet, rf, and Wilcoxon filter. The maximum number of selected features was set to 25 for all filters, with multiDEGGs features restricted to combined features only.
