## Supplementary figures and images for "multiDEGGs: a multi-omic differential network analysis package for biomarker discovery and predictive modeling"

### Supplementary Figure 3

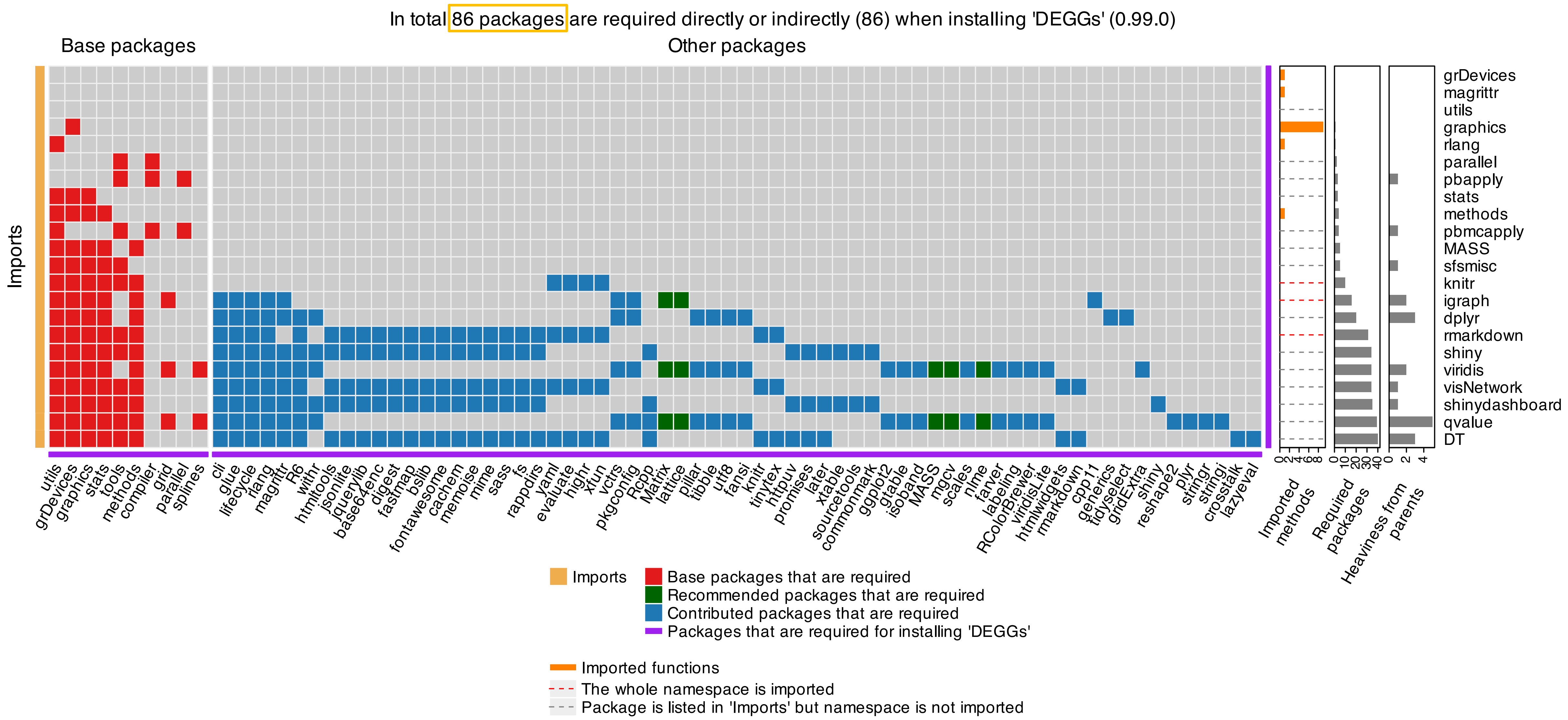


Supplementary Figure 3 – Heatmap of package dependencies for the DEGGs package.

### Supplementary Figure 4

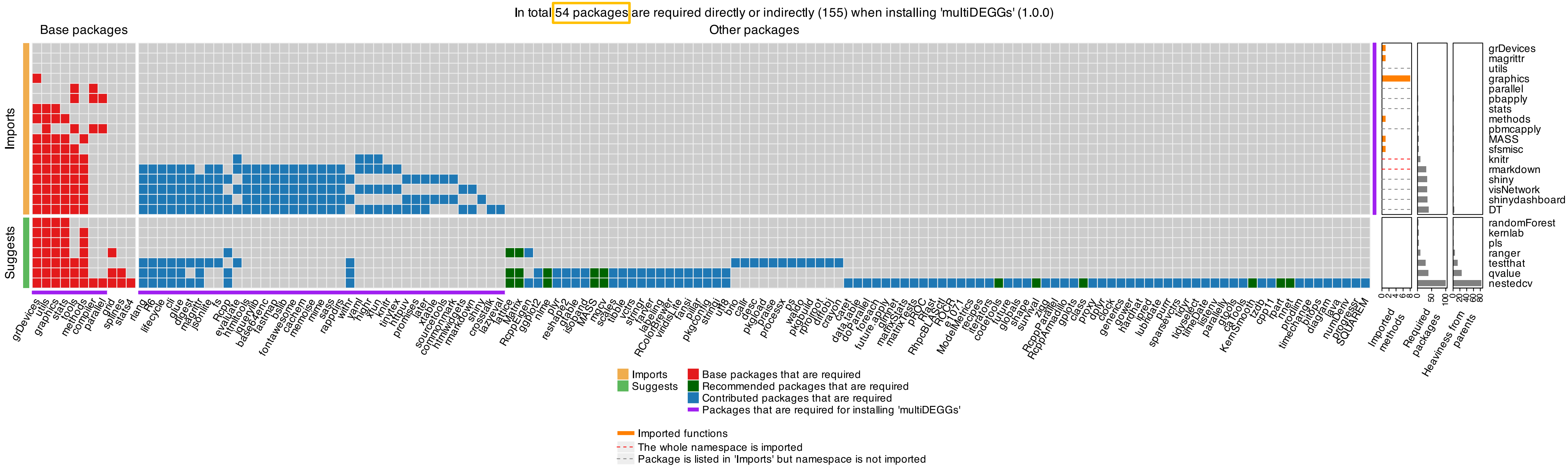


Supplementary Figure 4 – Heatmap of package dependencies for the multiDEGGs package.
